# Alternative pathway androgen biosynthesis and human fetal female virilization

**DOI:** 10.1101/609289

**Authors:** Nicole Reisch, Angela E. Taylor, Edson F. Nogueira, Daniel J. Asby, Vivek Dhir, Andrew Berry, Nils Krone, Richard J. Auchus, Cedric H.L. Shackleton, Neil A. Hanley, Wiebke Arlt

## Abstract

Androgen biosynthesis in the human fetus proceeds through the adrenal sex steroid precursor dehydroepiandrosterone, which is converted towards testosterone in the gonads, followed by further activation to 5α-dihydrotestosterone in genital skin, thereby facilitating male external genital differentiation. Congenital adrenal hyperplasia due to P450 oxidoreductase deficiency results in disrupted dehydroepiandrosterone biosynthesis, explaining undervirilization in affected boys. However, many affected girls are born virilized, despite low circulating androgens. We hypothesized that this is due to a prenatally active, alternative androgen biosynthesis pathway from 17α-hydroxyprogesterone to 5α-dihydrotestosterone, which bypasses dehydroepiandrosterone and testosterone, with increased activity in congenital adrenal hyperplasia variants associated with 17α-hydroxyprogesterone accumulation. Here we employ explant cultures of human fetal organs (adrenal, gonads, genital skin) from the major period of sexual differentiation and show that alternative pathway androgen biosynthesis is active in the fetus, as assessed by liquid chromatography-tandem mass spectrometry. We found androgen receptor expression in male and female genital skin using immunohistochemistry and demonstrated that both 5α-dihydrotestosterone and adrenal explant culture supernatant induce nuclear translocation of the androgen receptor in female genital skin primary cultures. Analyzing urinary steroid excretion by gas chromatography-mass spectrometry, we show that neonates with P450 oxidoreductase deficiency produce androgens through the alternative androgen pathway during the first weeks of life. We provide quantitative in vitro evidence that the corresponding P450 oxidoreductase mutations predominantly support alternative pathway androgen biosynthesis. These results indicate a key role of alternative pathway androgen biosynthesis in the prenatal virilization of girls affected by congenital adrenal hyperplasia due to P450 oxidoreductase deficiency.

**SIGNIFICANCE:** In the classic androgen biosynthesis pathway, testosterone is converted to 5α-dihydrotestosterone, a step crucially required for normal male genital virilization. Congenital adrenal hyperplasia (CAH) due to P450 oxidoreductase deficiency (PORD) is an inborn disorder that disrupts classic androgen biosynthesis. However, some affected girls present with severe genital virilization at birth. We hypothesized that this is explained by a prenatally active, alternative biosynthesis pathway to 5α-dihydrotestosterone. We show that adrenals and genital skin cooperate to produce androgens via the alternative pathway during the major period of human sexual differentiation and that neonates with PORD still produce alternative pathway androgens during the first weeks of life. This indicates that alternative pathway androgen biosynthesis drives prenatal virilization in CAH due to PORD.

## Introduction

Gonadal development depends on chromosomal sex, whereby the 46,XY or 46,XX karyotype, established at fertilization, dictates subsequent development of either testis or ovary (1–3). Gonadal hormones then direct differentiation of either male or female genitalia. In humans, sexual differentiation is established at 7-12 weeks *post conception* (wpc) (4).

While secretion of testosterone by fetal testis Leydig cells is thought sufficient to drive virilization of the internal genitalia in the male fetus (5), differentiation of the external genitalia requires the action of 5α-dihydrotestosterone (DHT), which is generated locally from circulating testosterone by the enzyme steroid 5α-reductase type 2 (SRD5A2) (6, 7). By contrast, differentiation of human female genitalia has been regarded as the default of a low androgen environment.

In humans, the regulation of sexual differentiation is intricately linked to early development of the adrenal cortex (4, 8). Disorders affecting adrenal steroidogenesis commonly affect sexual differentiation, as exemplified by the multiple variants of congenital adrenal hyperplasia (CAH), which result either in inappropriate or disrupted androgen biosynthesis. This consequently causes Disorders of Sex Development (DSD), which can manifest with external genital virilization in newborn girls (46,XX DSD), or undermasculinization of external genitalia in male neonates (46,XY DSD) (9). The most common variant of CAH, 21-hydroxylase (CYP21A2) deficiency, manifests with 46,XX DSD, while 17α-hydroxylase/17,20-lyase (CYP17A1) deficiency results in 46,XY DSD.

The congenital adrenal hyperplasia variant cytochrome P450 oxidoreductase (POR) deficiency can manifest with both 46,XY DSD and 46,XX DSD (10–12). POR plays a pivotal role as the obligatory electron donor to all microsomal cytochrome P450 enzymes, including CYP21A2 and CYP17A1, the latter catalyzing the biosynthesis of dehydroepiandrosterone (DHEA), the major precursor for testosterone biosynthesis. Consequently, POR deficiency (PORD) results in low circulating androgen concentrations, which readily accounts for 46,XY DSD, but fails to account for the severe virilization of external genitalia regularly observed in affected 46,XX neonates.

An explanation for this striking and seemingly contradictory genital phenotype in PORD has been lacking. We hypothesized that this apparent paradox could be explained by the existence of an alternative pathway to androgen production that generates DHT from 17α-hydroxyprogesterone (17OHP) during human fetal sexual differentiation, thereby bypassing the classic androgen biosynthesis pathway via DHEA and testosterone, as previously proposed by us (11) and others (13, 14). Elements of this pathway have been characterized in the fetal gonad of the tammar wallaby pouch young (15–17) and fetal opossum urogenital tract (18, 19). 17OHP accumulates in PORD but also in the most common CAH variant, 21-hydroxylase deficiency, and thus could feed into the proposed alternative pathway, if present in the fetus. Indirect biochemical evidence has indicated that the proposed alternative pathway is active postnatally in individuals with CAH due to 21-hydroxylase deficiency (20, 21) and may explain maternal virlization observed in pregnancies affected by PORD (22, 23). However, direct delineation of the putative alternative pathway during human fetal development, and in particular during the major period of sexual differentiation, has been lacking.

Here, we present conclusive evidence for the presence and activity of the alternative pathway in the human fetus, producing potent androgens during the major period of sexual differentiation, and we show that human fetal female external genitalia respond sensitively to androgens during the same period. In concert, these findings define an alternative pathway for androgen biosynthesis during the critical period of sexual differentiation in the human fetus that represents an important mechanism to explain the prenatal virilization of female infants affected by CAH.

## Results and Discussion

### Androgen biosynthesis in the human fetus during sexual differentiation

To ascertain the presence and activity of the hypothesized alternative androgen pathway, we performed incubations with male and female fetal adrenals, gonads and genital skin, which were collected at 6-10 wpc as previously described (8). We separately added deuterated steroid substrates for each step of the alternative pathway and employed liquid chromatography-tandem mass spectrometry (LC-MS/MS) to quantify the resulting products (**SI Appendix**, **Fig. S1** and **Tables S1+S2**). Experiments were conducted at least in triplicate for each organ of either sex (**SI Appendix**, **Figs. S2+S3**).

Explant incubations with female tissue showed that some 17OHP entered the classic androgen biosynthesis pathway, yielding androstenedione in adrenal, ovary and genital skin. However, testosterone was detected in only 2 of 9 female genital skin incubations, and not all in female adrenals and gonads (**Fig. 1A**). By contrast, all three female tissues (genital skin, ovaries, adrenals) showed conversion of 17OHP and subsequent intermediates through all steps of the proposed alternative androgen pathway, with the end product, DHT, produced from 5α-androsterone, with either 5α-androstanediol or also 5α-androstanedione as intermediates (**Fig. 1A**).

**Fig. 1:**
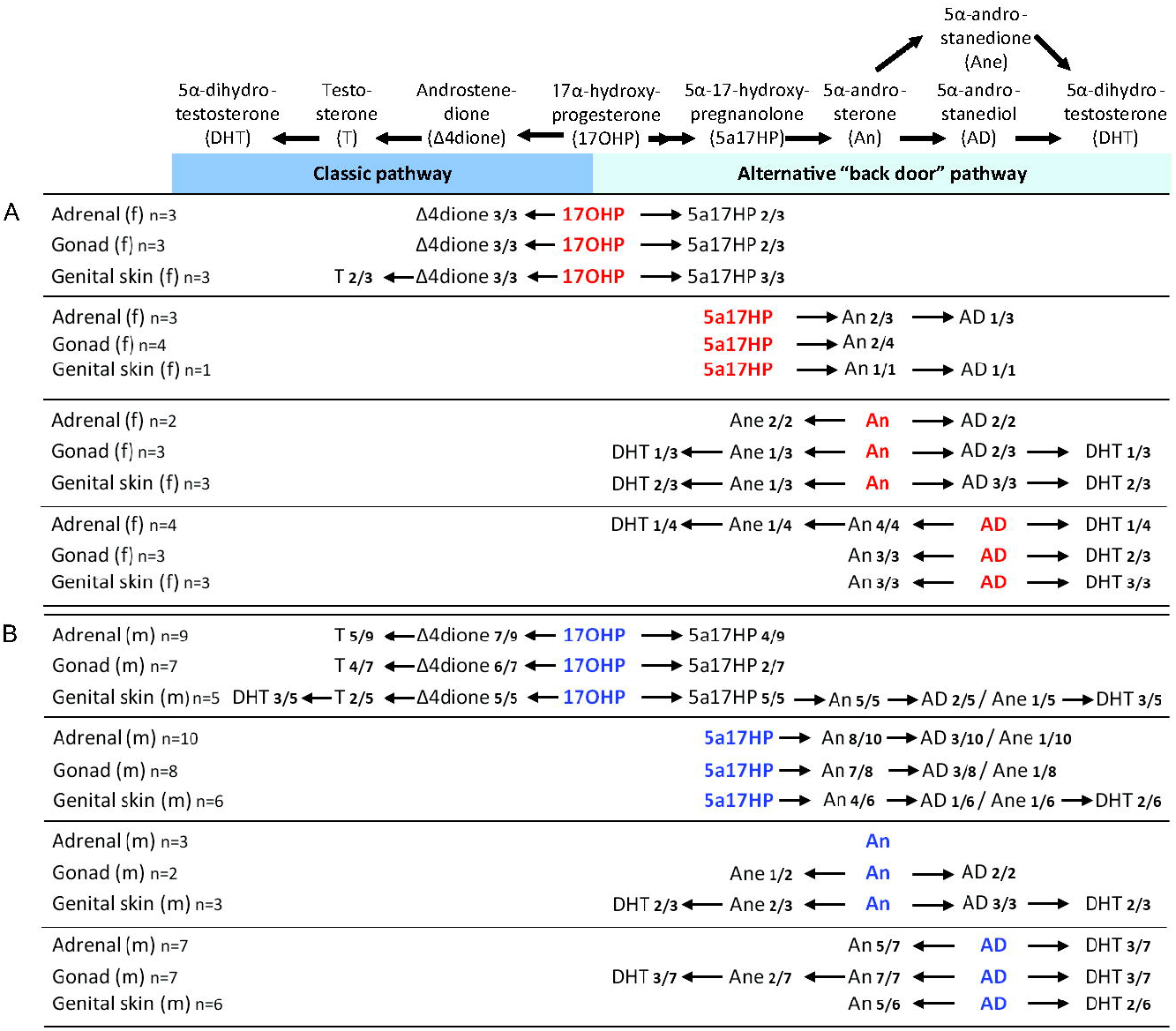
Androgen biosynthesis via classic and alternative androgen pathways as observed in human fetal organ explant cultures from the major period of sexual differentiation. The steroid substrates added to the explant cultures derived from fetal adrenals, gonads and genital skin (collected at 6-10 wpc) are shown in red and blue for fetal female (Panel A) and male (Panel B) tissues, respectively. Conversion products detected and identified by liquid chromatography-tandem mass spectrometry (LC-MS/MS) are given in black. f, female; m, male; n numbers indicate the number of biological replicates per tissue and fetal sex and how many of the cultures showed detectable synthesis of the indicated products.

Male fetal tissues readily converted 17OHP along the classic androgen pathway to testosterone and and, in the alternative pathway, to 5α-17-hydroxypregnanolone (5α-17HP). In male genital skin, conversion from 17OHP proceeded until the generation of DHT (**Fig. 1B)**, feasibly arising from either route, as DHT represents the end product of both classic and alternative androgen pathways. However, incubations with the intermediate substrates of the alternative pathway demonstrated stepwise catalysis to DHT by male adrenal, testis and genital skin (**Fig. 1B**). Consistent with studies in non-human species (24, 25), we did not identify 5α-pregnane-17α-ol-3,20-dione as a significant intermediate of the two-step conversion of 17OHP to 5α-17HP in tissues from either sex, likely due to its immediate onward conversion.

Taken together, these data show that adrenal, gonad and genital skin are capable of androgen biosynthesis via both classic and alternative pathway in both sexes. Our data suggest the presence of an integrated adreno-genital steroidogenic unit capable of producing DHT during the period of sexual differentiation in both male and female fetuses.

We did not study the recently described 11-oxygenated androgen biosynthesis pathway (26, 27), initiated by conversion of androstenedione to 11-hydroxyandrostenedione by CYP11B1 11β-hydroxylase activity, and eventually yielding 11-ketotestosterone, which activates the androgen receptor with similar potency to testosterone. Recent work indicated a significant role of 11-oxygenated androgens in CAH due to 21-hydroxylase deficiency (28) and polycystic ovary syndrome (29). We previously demonstrated presence and activity of CYP11B1 in the human fetal adrenal during the period of sexual differentiation (8), hence at least the initial step of the 11-oxygenated androgen pathway is likely to occur. However, urine steroid excretion analysis in infants with PORD (13) did not show increased excretion of 11-hydroxyandrosterone, the major metabolite derived from 11-oxygenated androgens, indicating that this side arm of the classic androgen pathway is unlikely to play a major role in PORD.

### Steroidogenic enzyme expression during human fetal sexual differentiation

The alternative pathway conversion of 17OHP to 5α-17HP requires sequential 5α-reductase and 3α-hydroxysteroid dehydrogenase activities.

Of the two isoforms, the first step is expected to require catalysis by steroid 5α-reductase type 1 (SRD5A1), since SRD5A2 does not to convert 17OHP efficiently (30). A study of fetal tissues from 12-20 wpc did not detect SRD5A1 in adrenals, gonads or genital skin (31); a recent study also using fetal tissues from the second trimester of pregnancy (11-21 wpc) described detection of SRD5A1 in liver, placenta, testis and genital tubercle, but not in the adrenal, while female gonads were not studied (32) However, studying tissue from the major period of human sexual differentiation, we found mRNA expression of SRD5A1 in adrenal, gonads and genital skin of both sexes (**Fig. 2A**). The second step, the conversion of 5α-pregnane-17α-ol-3,20-dione to 5α-17HP requires 3α-hydroxysteroid dehydrogenase activity, in keeping with the observed expression of *AKR1C1* and *AKR1C3*, both of which encode enzymes capable of catalyzing this reaction (**Fig. 2A**).

**Fig. 2:**
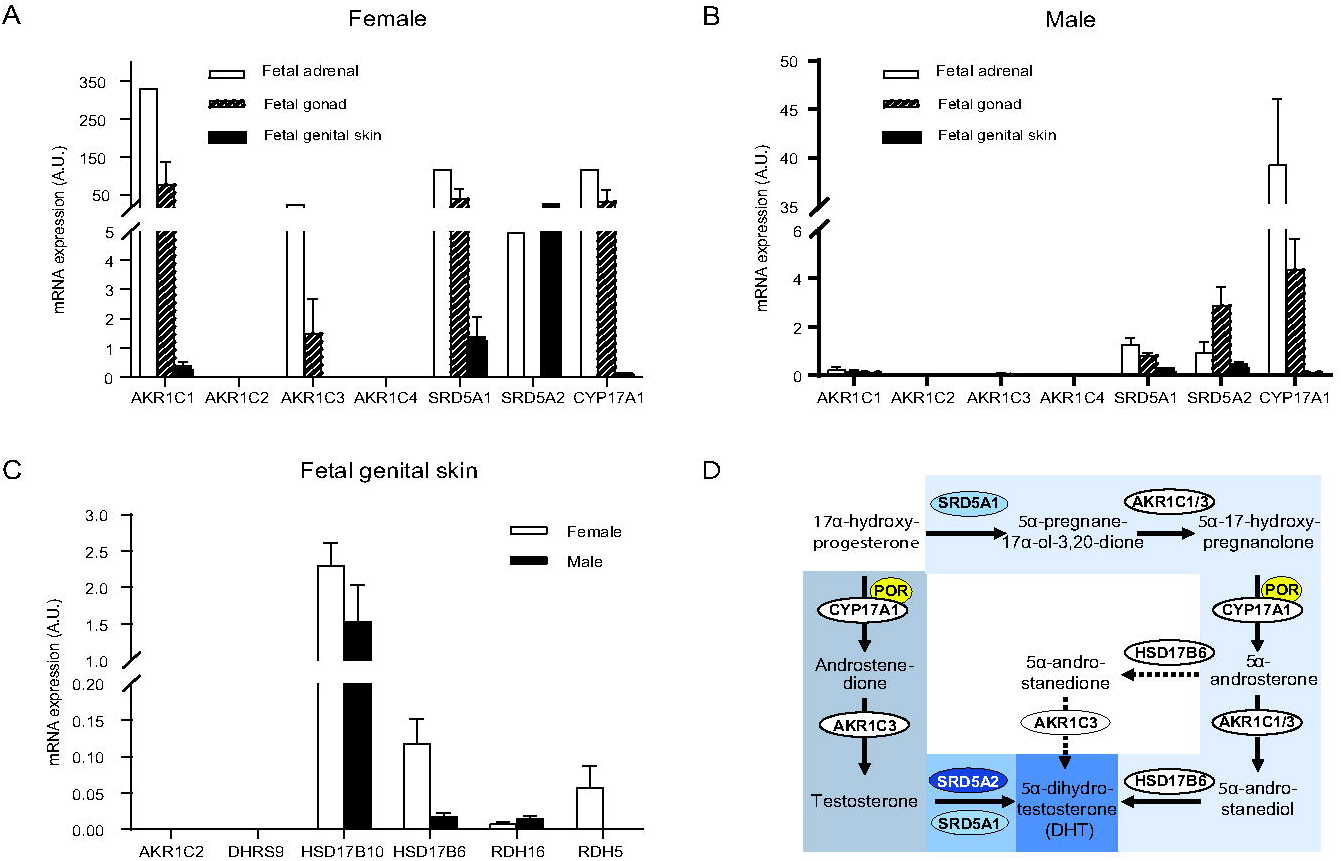
Steroidogenic enzyme expression in human fetal tissue from the major period of sexual differentiation and their proposed role(s) in alternative pathway synthesis. Panels A+B, mRNA expression (mean+SEM) in human fetal tissues collected at 6-10 wpc as measured in by qPCR in female adrenal (n=1), gonad (n=2), genital skin (n=2) and corresponding male tissues (≥3 biological replicates for adrenal, gonads, and genital skin). Expression data were normalized to ribosomal 18S. Panel C, Schematic summary of the proposed distinct roles of the identified enzymes in the classic androgen pathway (dark blue) and the alternative androgen synthesis pathway (light blue) both resulting in the synthesis of potent 5α-dihydrotestosterone. Arrows indicate observed conversions in the fetal organ explant cultures; dotted arrows represent reactions only rarely observed.

The next step in the alternative pathway requires CYP17A1 17,20 lyase activity; 5α-17HP is the preferred substrate for this reaction and efficiently converted to 5α-androsterone (33). We detected robust *CYP17A1* expression in human fetal adrenals from the major period of sexual differentiation, consistent with previous reports (8, 34) and also in the gonads of both sexes (**Fig. 2B**). The subsequent reduction of 5α-androsterone to 5α-androstanediol requires 17β-hydroxysteroid dehydrogenase activity, which can be provided by AKR1C3 or AKR1C1, expressed in adrenal, gonads, and genital skin (**Fig. 2A**).

The final step of the proposed pathway involves the conversion of 5α-androstanediol to DHT, which requires 3β-epimerase (oxidative 3α-HSD) activity. Several enzymes have been considered to catalyze this reaction (i.e. AKR1C2, RDH5, DHRS9, HSD17B10, HSD17B6 and RDH16). However, only HSD17B6 and RDH16 are capable of efficient oxidation of 5α-androstanediol to DHT, as previously demonstrated by transactivation of the androgen receptor following cell-based overexpression (24, 35). We found expression of both HSD17B6 and RDH16 in fetal genital skin of both sexes (**Fig. 2C**).

In summary, we detected the transcripts encoding all enzymes required to catalyze the alternative androgen pathway (**Fig. 2D**). Taken together with the steroid conversion studies, these data comprehensively demonstrate that the normal adrenal, gonad and genital skin are capable of androgen biosynthesis via both classic and alternative pathway in both sexes. Our data point to an adreno-genital steroidogenic unit that can cooperate to produce DHT via the alternative pathway during the major period of human sexual differentiation.

### Androgen receptor in female genitalia from the start of sexual differentiation

Having demonstrated the capacity for DHT production from steroidogenic precursors in female fetuses, we corroborated its ability to function by examining the presence of the androgen receptor (AR) in female external genitalia from the start of sexual differentiation. Previously, AR expression was documented in four female human fetuses from 9-18 wpc (36). Inour study, we readily detected AR protein in stromal cells in the urethral folds of the external genitalia in both male and female fetuses at the onset of sexual differentiation (**Fig. 3A-C**). Its nuclear localization in fixed tissues in both sexes implied AR was ligand-bound. To explore this further we studied the intracellular localization of AR by immunofluorescence in female external genitalia fibroblasts taken into primary culture from the same stage of development. In steroid-free media, the external genitalia cells demonstrated cytoplasmic AR localization. As expected, the addition of 1 nM DHT induced nuclear translocation of AR. Strikingly, the same translocation was observed when using medium conditioned overnight from the corresponding female adrenal gland (**Fig. 3D**). In combination, these data show AR from the start of sexual differentiation in both male and female external genitalia and demonstrate a functional adreno-genital steroidogenic unit capable of causing AR nuclear translocation.

**Fig. 3:**
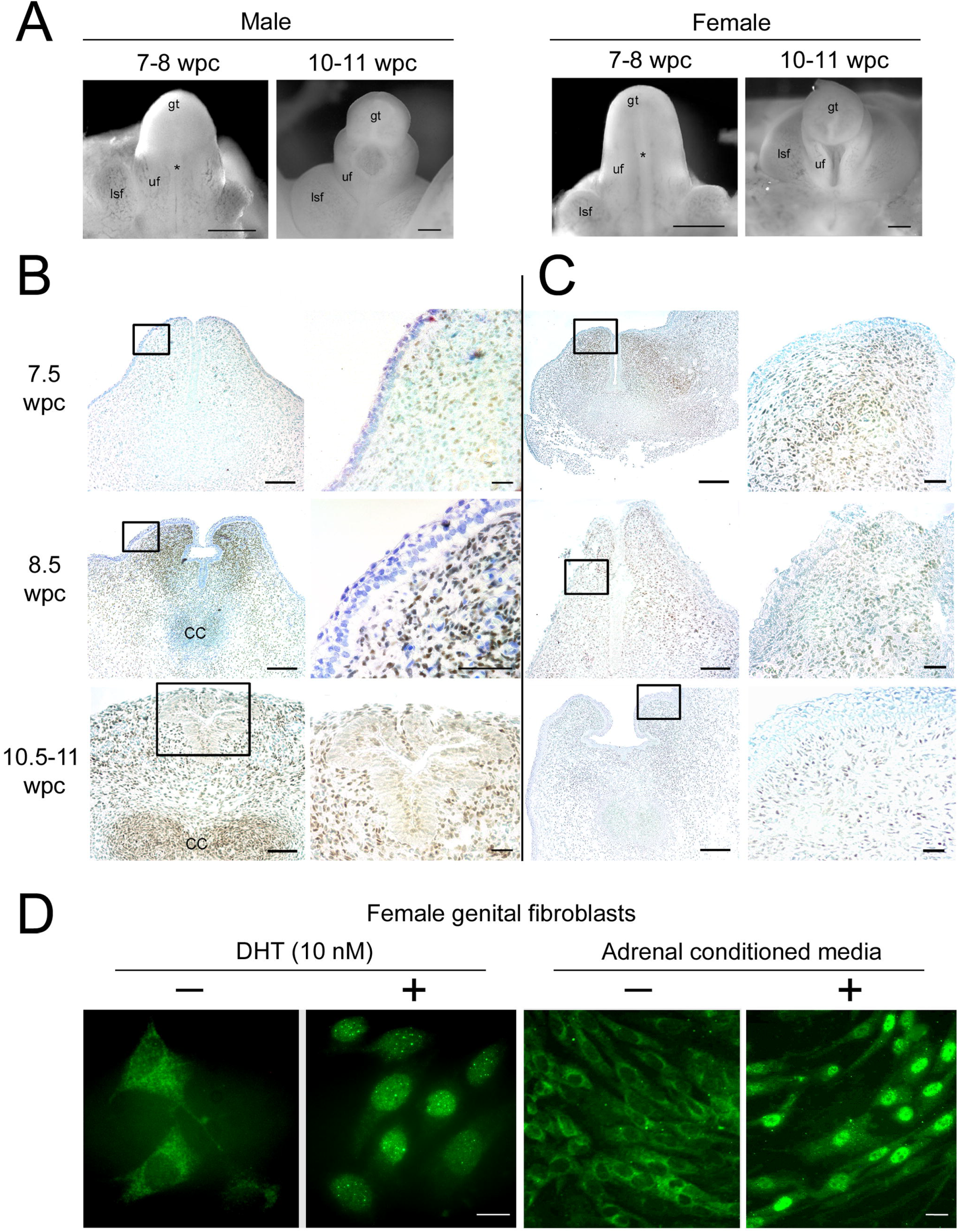
The androgen receptor is present in both male and female genitalia from the onset of sex differentiation. Panel A, Morphology of male and female external genitalia at the onset of sex differentiation. gt, genital tubercle; uf, urethral fold; lsf, labioscrotal fold; *, patency between urethral folds at 7-8 wpc that is partially sealed in males by 10-11 wpc. Panels B+C, Immunohistochemistry in trasverse sections through the phallus for AR in male (Panel B) and female (Panel C) external genitalia from the start of human sexual differentiation at 7-8 wpc counter-stained with toluidine blue. Boxes in the left-hand panel are shown at higher magnification to the right. Panel D, Immunofluorescence for AR in female external genitalia fibroblasts in the presence (+) or absence (−) of 10nM DHT (left) or medium conditioned by overnight incubation with an adrenal gland from the same female fetus (right). Size bars: Panel A, 500μm; Panels B+C, 100μm (low) and 20μm (high magnification); Panel D, 25μm.

### Androgen biosynthesis in neonates with congenital adrenal hyperplasia due to P450 oxidoreductase deficiency

Having shown evidence of the alternative pathway during human sexual differentiation, we next investigated whether we could demonstrate equivalent activity *in vivo*. To address our hypothesis that excess alternative pathway androgen biosynthesis in fetal life explains female virilization (46,XX DSD) in CAH due to PORD (12, 37), in whom the classic androgen pathway is disrupted, we identified three patients with a mutation known to cause 46,XX DSD in affected girls and facilitate normal male appearance of external genitalia in affected boys (POR A287P) (11, 12). All three individuals had a 46,XY karyotype; patients 1 and 2 harbored homozygous A287P mutations while patient 3 was compound heterozygous for POR A287P/G188_V191dup. While postnatal circulating androgens in PORD are low in infancy and beyond (12, 37), we hypothesized that affected individuals would still show evidence of excess alternative pathway androgen biosynthesis in the immediate neonatal period. We analysed urinary steroid metabolite excretion by gas chromatography-mass spectrometry and detected significant alternative pathway activity in the neonatal period, consistent with previous preliminary findings that argued for the adrenal gland as a major site for alternative pathway activity (13). During the first three weeks after birth we documented increased excretion of the 17OHP metabolite 17-HP compared to healthy controls (**Fig. 4A**), consistent with accumulation of 17OHP, the initial substrate of the alternative androgen pathway. We found significantly increased excretion of 5α-17HP **(Fig. 4B**), a key intermediate of the alternative pathway; and finally, there was significantly increased excretion of the major DHT metabolite 5α-androsterone (**Fig. 4C**). By contrast, etiocholanolone excretion, derived from classic pathway androgen biosynthesis, did not differ between the affected and healthy control groups (**Fig. 4D**). This result corroborates a previous report on urine steroid metabolite excretion in a 46,XY infant affected by PORD, which also described increased 5α-androsterone but unremarkable etiochaolanolone excretion (38). Of note, the POR A287P homozygous patients in our study had higher excretion of 5α-17HP and 5α-androsterone than the compound heterozygous patient, who can be expected to have less alternative pathway activity due to the greater impairment of residual POR function. Taken together, these findings indicate increased *in vivo* alternative pathway androgen biosynthesis in neonates with PORD, which ceases shortly after birth.

**Fig. 4:**
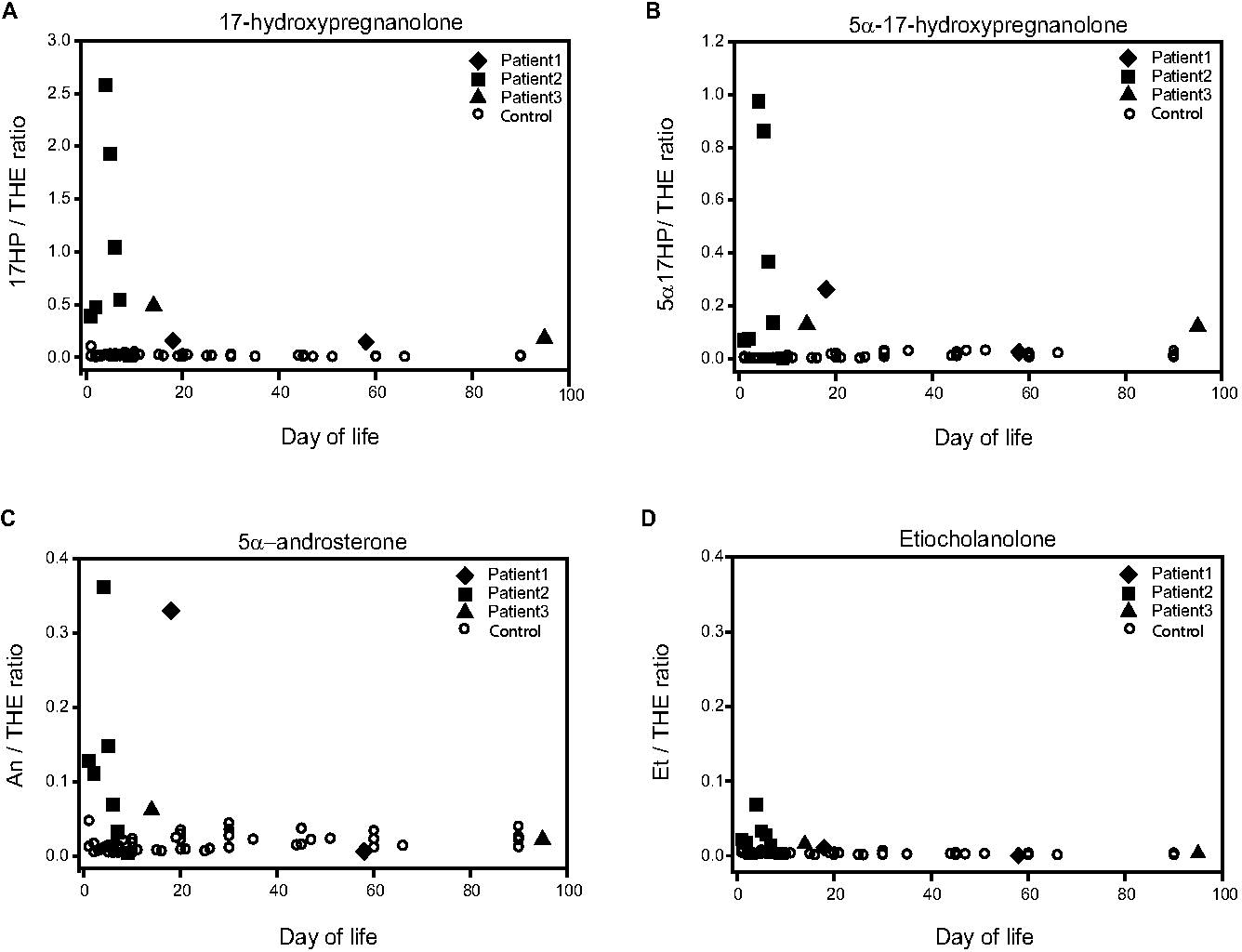
Urinary steroid excretion in three 46,XY neonates with POR deficiency (closed symbols) in comparison to nine sex- and age-matched healthy controls (open symbols). POR deficient neonates harbored the A287P mutation, which in the homozygous state is associated with normal male genitalia in boys and genital virilization (46,XX DSD); we included data from three 46,XY neonates with PORD; two harbored homozygous POR mutations (A287P/A287P), one compound heterozygous mutations (A287P/G188_V191dup). Longitudinal urine collections were carried out during the first three months of life and analyzed by gas chromatography-mass spectrometry. Depicted are the urinary excretion of (Panel A) the 17OHP metabolite 17-hydroxypregnanolone (17HP) and the two alternative pathway intermediates (Panel B) 5α-17-hydroxypregnanolone (5α-17HP) and (Panel C) 5α-androsterone (An), in comparison to (Panel D) etiocholanolone (Et), which is only generated via the classic androgen pathway. All steroids are shown relative to tetrahydrocortisone (THE), an abundant adrenal-derived steroid metabolite, as the denominator.

These findings were further supported by in vitro 17,20 lyase activity assays employing yeast microsomes co-transformed with human CYP17A1 and either wild-type or mutant POR. The results demonstrated that the residual activity of POR A287P was higher in the alternative androgen pathway than in the classic pathway. POR A287P demonstrated significantly higher activity than POR mutant H628P, which in the homozygous state is associated with severe male undervirilisation (46,XY DSD) and normal female genital phenotype (**SI Appendix**, **Fig. S1A-C**). Additional experiments with yeast microsomes co-transformed with CYP19A1 and wild-type or mutant POR demonstrated that neither mutant affected aromatase activity (**SI Appendix**, **Fig. S1D**), thus excluding impaired aromatization of classic pathway androgens as a driver of prenatal virilization.

In conclusion, we have provided in vitro, ex vivo, and in vivo evidence for the existence and activity of an alternative pathway for the synthesis of the most potent androgen, DHT, during early human development. Our data demonstrate that, through cooperation of an adreno-genital steroidogenic unit, the alternative androgen pathway yields active androgen synthesis in the female fetus, with excess activity driving female virilization, 46,XX DSD, in CAH due to P450 oxidoreductase deficiency. Given that the alternative pathway substrate 17OHP also accumulates in 21-hydroxylase deficiency, it is conceivable that alternative pathway androgens contributes to prenatal virilization in this most common CAH variant.

## MATERIALS AND METHODS

### Collection of human embryonic and fetal material

Ethical approval for these studies was granted by the North West Haydock Research Ethics Committee of the UK Health Research Authority (approval number 18/NW/0096). The collection and staging of human embryonic and fetal material was carried out with informed consent, as described previously (8, 39), using the Carnegie classification and fetal foot length to provide a direct assessment of developmental age as days or weeks post conception (dpc or wpc), respectively, and male fetal material was identified by SRY expression, as previously described (1). We analyzed organs and tissue from 30 fetuses: 25 male and 5 female; median age 55 dpc (range 44-84 dpc).

### RNA extraction, reverse transcription, and quantitative PCR

Total RNA was extracted from whole organs using the Tri-Reagent system (Sigma-Aldrich). RNA integrity and concentrations were assessed using the Nanodrop spectrophotometer (Wilmington, DE). Reverse transcription was carried out employing a standard protocol. mRNA expression levels were quantified using an ABI 7500 sequence detection system (Perkin-Elmer Applied Biosystems, Warrington, U.K.), employing Applied Biosystems “assay on demand” probe and primers for specific amplification of SRD5A1, SRD5A2, CYP17A1, AKR1C1, AKR1C2, AKR1C3, AKR1C4, HADH2/HSD17B10, HSD17B6, RDH5, DHRS9 and RDH16 (for further details see **SI Appendix**).

### Tissue explant culture and steroid identification by tandem mass spectrometry

Whole organ tissue explants (fetal adrenals, gonads, and genital skin) were cultured in DMEM/F12 (PAA Laboratories Inc.) supplemented with 2% Ultroser SF (i.e. steroid free; BioSepra) and 1% ITS+ (BD) at 37°C in humidified 5% CO_2_ and 95% air for 64 hours. Genital skin was cultured as monolayers and used for experiments at passage four.

Identification of steroid products from the explant cultures was achieved using liquid chromatography-tandem mass spectrometry (LC-MS/MS). Steroids were positively identified by comparison of retention time and MS/MS mass transitions to authentic steroid standards (**SI Appendix**, **Table S1**). Two mass transitions were used to positively identify each steroid referred to as quantifier and qualifier ions, respectively; the resolution of a series of authentic steroid standards is shown in the **SI Appendix**, **Fig. S2**, alongside further method details.

For steroid conversion assays, tissue explants were incubated with precursor steroids purchased from Steraloids, Inc. (Newport, RI) and Sigma-Aldrich (Dorset, UK). For explant cultures assessing the conversion of 17-hydroxyprogesterone, 5α-androsterone, and 5α-androstanediol, we used deuterated steroids (for details see **SI Appendix**, **Table S2**). Representative results of steroid detection following explant culture incubations are shown in the **SI Appendix**, **Figs. S3+S4**, for female and male tissues.

### Immunohistochemistry

Immunohistochemistry, immunoblotting and immunofluorescence were carried out as reported previously (40), using monoclonal mouse anti-AR (1:100, Lab Vision Corp, Cheshire, UK).

### Urine steroid metabolite excretion analysis

Urine samples were collected longitudinally from birth until 90 days of life in three neonates affected with PORD (two homozygous for POR A287P, the other one compound heterozygous for POR A287P/G188_V191dup) and compared to those collected from nine healthy controls. These were matched for sex, age and gestational age at birth and collected during the same time window. The parents of PORD patients and healthy controls provided written informed consent prior to urine collection. The study protocol was reviewed and approved by the Research Ethics Committee of University College London Institute of Child Health/Great Ormond Street Hospital NHS Trust (REC reference 05/Q0508/24).

Urinary steroid hormone profiles were determined by gas chromatography-mass spectrometry analysis as described previously (41). The final analytical samples are the methyloxime-trimethylsilyl (MO-TMS) derivatives of steroids enzymatically released from sulfate and glucuronide conjugation. Analytes quantified by selected ion monitoring were normalized to tetrahydrocortisone (THE), the most abundant steroid metabolite consistently excreted throughout life, with no significant difference in urinary THE concentrations identified between PORD patients (n=3; 13 urine samples; median 285 μg/L, range 56-1256 μg/L) and healthy controls (n=9; 48 urine samples; median 306 μg/L, range 59-1663 μg/L).

## Supporting information

Suppl. Appendix incl. Suppl Figs 1-4 and Suppl tables 1-2

## ACKNOWLEDGMENTS

This work was supported by the Wellcome Trust (Senior Research Fellowship WT088566, to N.A.H.; Investigator Award in Science 209492/Z/17/Z, to W.A.), the Medical Research Council UK (Program Grant 0900567, to W.A.), the Gerald Kerkut Trust (PhD studentship to D.J.A.), the European Commission (Marie Curie Intra-European Fellowship PIEF-GA-2008-221058, to N.R.), the Deutsche Forschungemeinschaft (Heisenberg Professorship 325768017, CRC/Transregio 205/1 ‘The Adrenal: Central Relay in Health and Disease’. to N.R.), and the Society for Endocrinology UK (Lab visit award, to N.R.). W.A. receives support from the National Institute for Health Research (NIHR) Birmingham Biomedical Research Centre at the University Hospitals Birmingham NHS Foundation Trust and the University of Birmingham (Grant Reference Number BRC-1215-20009). The views expressed are those of the authors and not necessarily those of the NIHR or the Department of Health and Social Care UK.

We thank Beverly A. Hughes, University of Birmingham, for technical support. We thank our research nurses and study participants for tissue collection.

