## Supplementary material for "Alternative pathway androgen biosynthesis and human fetal female virilization": Suppl. Appendix incl. Suppl Figs 1-4 and Suppl tables 1-2

**This Supplemental File includes:**

Supplementary Results

Supplementary Methods

Supplementary References

Figures S1 to S4

Tables S1 to S2

### Supplementary Results

#### *Androgen biosynthesis in neonates with CAH due to P450 oxidoreductase deficiency*

We were able to longitudinally collect spot urines in three neonates affected by PORD. All three neonates had a 46,XY karyotype. We previously reported genotype and phenotype of the three PORD neonates as part of a larger PORD cohort (1); in that paper, patients 1, 2 and 3 were reported as patients 8, 16, and 17, respectively); below we have briefly summarized information on genotype and the key clinical characteristics they presented with at birth.

Patient 1 (46,XY; POR A287P/A287P) was born at 34 weeks of gestation of a twin pregnancy; his clinically unaffected twin sister (46,XX) is heterozygous for POR A287P. At birth, his external genitalia had normal male appearance, with bilaterally descended testes.

Patient 2 (46,XY; POR A287P/A287P) was born at 35 weeks gestation and is the younger sibling of a girl who had been born with severe virilization (46,XX DSD). Patient 2 presented with normal male genitalia including bilaterally descended testes.

Patient 3 (46,XY; POR A287P/G188_V191dup) was born at full term. During the second and third trimester the mother had reported the development of facial acne and slight coarsening of facial features, which both quickly resolved after birth. At birth, there were overt signs of undervirilization (46,XY DSD) with a small midline phallus measuring 1.5cm, a bifid scrotum with palpable testes and perineoscrotal hypospadias.

As patient 3 is compound heterozygous for the major loss-of-function mutant G188_V191dup, his combined residual POR activity can be assumed to be lower than in patients 1 and 2, who are compound heterozygous for POR A287P, a missense mutation previously shown to have considerable residual activity. This difference in residual POR activity is also supported by their malformation phenotypes, which occurs in PORD most likely due to impaired function of CYP enzymes involved in sterol biosynthesis and retinoic acid metabolism. While the two boys homozygous for POR A287P presented with only mild to moderate malformations, patient 3 had a more severe malformation phenotype (1).

Interestingly, maternal virilization was only observed during the pregnancy with patient 3 and not in the two boys with homozygous POR A287P; maternal virilization was only observed in one of five previously reported pregnancies affected by homozygous POR A287P mutations (2). As POR A287P has previously been shown not to impair CYP19A1 aromatase activity (3), it is tempting to assume that impaired placental aromatase activity is an important contributor to maternal virilization, as also previously proposed by others (4, 5).

***Differential effects of POR mutants on sex steroid biosynthesis***

Both the classic and the alternative pathway to androgens require the 17,20-lyase activity of the steroidogenic enzyme CYP17A1. In the classic pathway, CYP17A1 converts 17-hydroxypregnenolone to DHEA and can also convert 17-hydroxyprogesterone (17OHP) to androstenedione (**Fig. S1A**). However, in humans, the latter reaction occurs with significantly lower efficacy (6) and, thus, only provides a relevant contribution to androgen synthesis if 17OHP accumulates, a pathognomonic finding in congenital adrenal hyperplasia with impaired 21-hydroxylase activity, typically due to inactivating *CYP21A2* mutations.

The proposed alternative androgen synthesis pathway starts with 17OHP and requires CYP17A1 17,20 lyase activity further downstream, to convert 5α-17-hydroxyallopregnanolone (5α-17HP) to androsterone. That reaction is strongly preferred by the human CYP17A1 enzyme (7), which means if 17,20 lyase activity is partially compromised, alternative pathway activity is the most likely one to still occur.

In congenital adrenal hyperplasia due to POR deficiency, the relationship between genotype and genital phenotype has a broad spectrum and its assessment is complicated by the fact that the majority of affected individuals are compound heterozygous (1, 4, 8). In our previously published cohort of 30 patients (1) we identified six individuals homozygous for POR A287P, four with 46,XX DSD whereas the two 46,XY individuals had normal male external genitalia. We recently identified two sisters homozygous for POR H628P, previously only described in the compound heterozygous state. The older sister had presented with primary amenorrhea and unambiguously female external genital appearance; chromosome analysis confirmed 46,XY DSD. Her younger sister had a 46,XX karyotype and normal female genitalia. Thus, the genital phenotypes relating to these two homozygous POR mutations represent exact mirror images, with 46,XX DSD for A287P and 46,XY DSD for H628P, respectively; however, in all affected individuals postnatal circulating androgens were found to be low. We hypothesized that this difference might be explained by a differential impact of the two mutations on CYP17A1 17,20 lyase activity within the classic and alternative androgen pathways during human fetal development.

To test this, we employed yeast microsomes co-expressing human CYP17A1 with human wild-type and mutant POR, respectively, and incubated them with the substrates for the classic and alternative androgen biosynthesis pathways, respectively, 17-hydroxypregnenolone (17-Preg) and 5α-17-hydroxypregnanolone (5α-17HP). Comparison of the two POR mutants revealed that A287P retained significantly higher activity than H628P in both the classic and the alternative androgen pathway conversions (**Fig. S1A+B**). POR A287P demonstrated higher activity in the alternative than in the classic androgen pathway, providing quantitative evidence supporting a more significant contribution of the alternative androgen pathway in conveying prenatal virilization in 46,XX individuals homozygous for POR A287P. CYP17A1 co-expressed with wild-type POR also exhibited slightly higher activity in the alternative vs. the classic androgen pathway conversion (**Fig. S1A+B**), consistent with the previous report describing 5α-17HP as the preferred CYP17A1 substrate (7).

The enzyme P450 aromatase (CYP19A1) also requires POR as an electron donor and patients with inactivating CYP19A1 mutations develop androgen excess due to decreased conversion of androgens to estrogens (9). Thus, it is theoretically possible that reduced androgen synthesis via the classic pathway could still result in androgen excess, dependent on the distinct effect of a specific POR mutant on CYP19A1 activity, as suggested previously (3, 4, 10). To exclude a role of impaired aromatization in the observed genital phenotype differences, we also examined the influence of POR mutants A287P and H628P on CYP19A1 activity, again employing a humanized yeast microsomal co-expression assay. Both POR mutants showed similar aromatase activity to wild-type POR for the conversion of androstenedione to estrone, when co-expressed with CYP19A1. These findings exclude impaired aromatization as the cause of the observed genital phenotype differences (**Fig. S1C**), also corroborating a previous report demonstrating normal aromatase activity in the presence of POR A287P (3).

### Supplementary Methods

#### *Steroid analysis in tissue explant culture supernatant by tandem mass spectrometry*

Steroids were extracted from 500µL of cell media by liquid-liquid extraction using 5mL of dichloromethane. The dichloromethane was evaporated under nitrogen at 55˚C and the steroids reconstituted in 100µL of 50/50 methanol/water. LC-MS/MS was carried out employing a Xevo TQ mass spectrometer combined with an Acquity uPLC system with an electro-spray ionisation source in positive ion mode (Waters, Elstree). Steroids were eluted from a BEH $C_{18}$ 2.1 x 50mm 1.7µm column using a methanol/water gradient system. Solvent A was 0.1% formic acid in water, and B was 0.1% formic acid in methanol. The flow rate was 0.6 mL/min, and starting conditions were 45% B increasing linearly to 75% B over 5 minutes.

#### *Quantitative PCR*

Reactions were performed in 10 µl volumes on 96-well plates in reaction buffer containing 2x TaqMan Universal PCR Master mix (Applied Biosystems, Foster City, CA). All reactions were normalized against 18S rRNA as an internal reference gene. Data were obtained as Ct values (Ct = cycle number at which logarithmic PCR plots cross a calculated threshold line) and used to determine Δct values with Δct = (ct of the target gene) – (ct of the housekeeping gene). Data are expressed as arbitrary units using the following transformation (expression $=10^5 \times [2^{-\Delta ct}]$ arbitrary units [AUs]).

#### *In vitro functional studies with humanized yeast microsomes*

Saccharomyces cerevisiae strain W303B was transformed with V10-CYP17A1 and V10-CYP19A1, respectively, using a modification of the lithium acetate method as previously described (11). Wild-type human CYP19A1 full-length cDNA was amplified from pcDNA3.1/hsCYP19A1 (generous gift from Evan Simpson, Prince Henry's Institute of Medical Research, Melbourne, Australia) using the 5' sense primer CAAAGATCTGGAACACAAGATGGTTTTGGAAA and the 3' antisense primer CCAAGATCTGCCTTCTCTAGTGTTCCAGACAC that both included BglII restriction sites for subsequent cloning into yeast expression vector V10.

Yeast stably expressing wild-type human CYP17A1 or CYP19A1 were then transformed with yeast expression vector pYcDE2, containing the coding sequence of either wild-type or mutant (A287P or H628P) human P450 oxidoreductase (POR). Yeast microsomes co-expressing POR with CYP17A1 and CYP19A1, respectively, were prepared as previously described (11). Microsomal enzyme activity assays were carried out with 0.5-5.0 µM [$^3$H]-17-hydroxypregnenolone or [$^3$H]-5α-17-hydroxypregnanolone for assessment of CYP17A1 17,20-lyase activity in the classic and the alternative androgen pathway, respectively, and with 10-500nM [$^3$H]-androstenedione for CYP19A1 aromatase activity assays. [$^3$H]-androstenedione (55.4 Ci/mol) was purchased from Amersham Biosciences (Amersham, UK) and [$^3$H]-pregnenolone from

PerkinElmer Life Sciences, Inc. (Boston, MA, USA); 17-Preg and 5α-17HP were custom synthesized as previously described (11, 12).

All reactions were started by the addition of 10mM NADPH to the microsomal incubation; assays assessing 17,20 lyase activity also contained 10pM of recombinant human cytochrome b5 (Invitrogen, Paisley, UK). All assays were carried out within the linear time range of the enzymatic reaction, as established by preceding experiments. Steroids were extracted using dichloromethane and separated by thin layer chromatography on PE SIL G/UV silica gel plates (Whatman, Maidstone, Kent, UK) using a 3:1 chloroform/ethyl acetate solvent system for CYP17A1 and 92.5:7.5 dichlormethane:acetone for CYP19A1. TLC scanner analysis (Bioscan 2000 image analyzer, Lablogic, Sheffield, UK) was employed for quantification of steroid substrates and products. All data represent the results of three independent triplicate experiments and are expressed as mean±SEM.

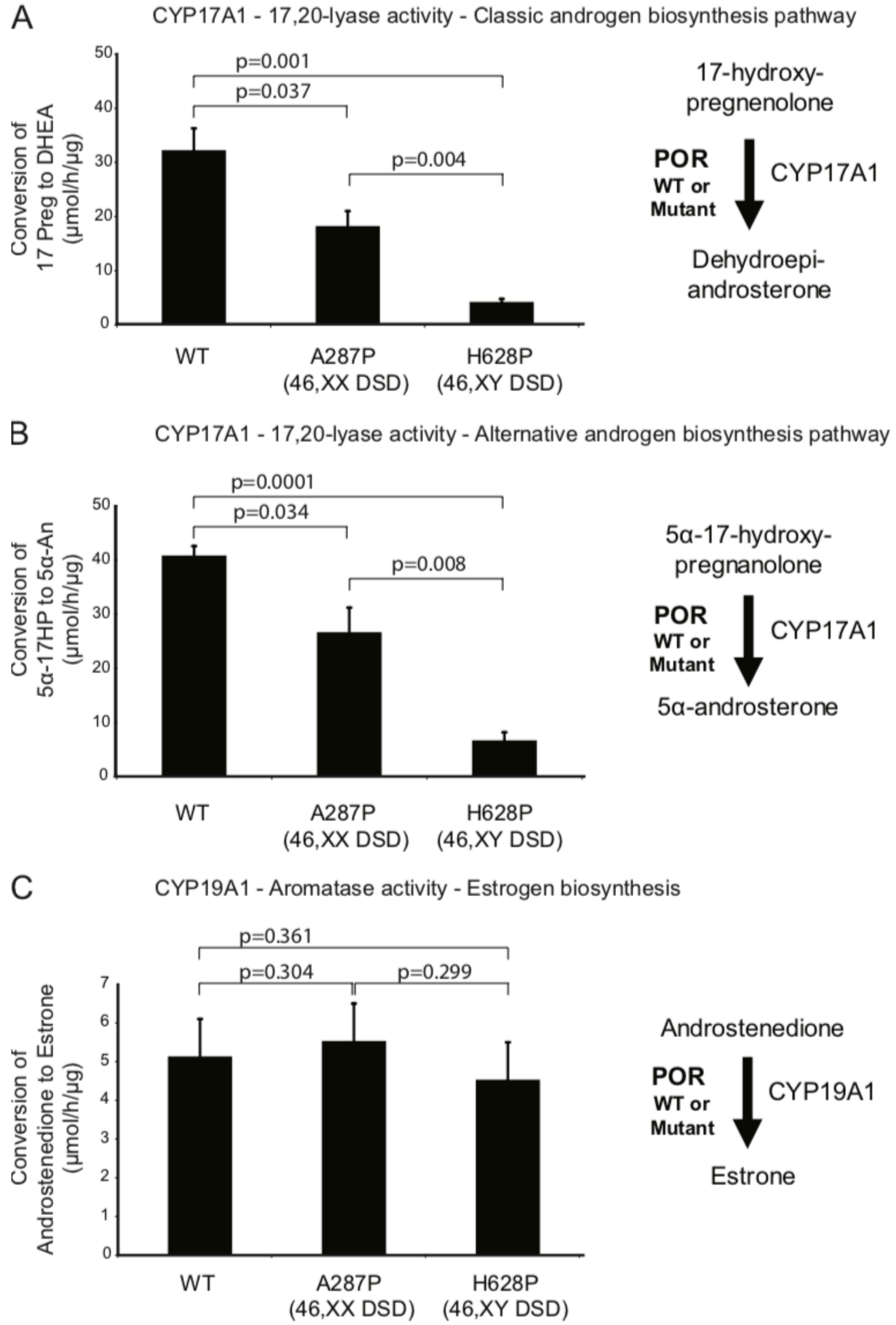


**Fig. S1. Differential effects of the POR mutants A287P and H628P, associated with 46,XX DSD and 46,XY DSD, respectively, on human sex steroid biosynthesis**. Conversion assays were carried out with yeast microsomes co-transformed with human CYP17A1 and wild-type or mutant POR, assessing conversions catalysed by CYP17A1 17,20 lyase activity: 17-hydroxypregnenolone to DHEA in the classic androgen pathway (**Panel A**) and 5α-17-hydroxypregnanolone to 5α-androsterone in the alternative androgen pathway (**Panel B**). **Panel C** shows CYP19A1 aromatase activity assays, employing yeast microsomes co-transformed with CYP19A1 and POR, assessing the conversion of androstenedione to estrone. All assays were carried out in three independent triplicate experiments and within the linear time range of the respective enzymatic reaction, as established by preceding experiments. Data are expressed as mean±SEM.

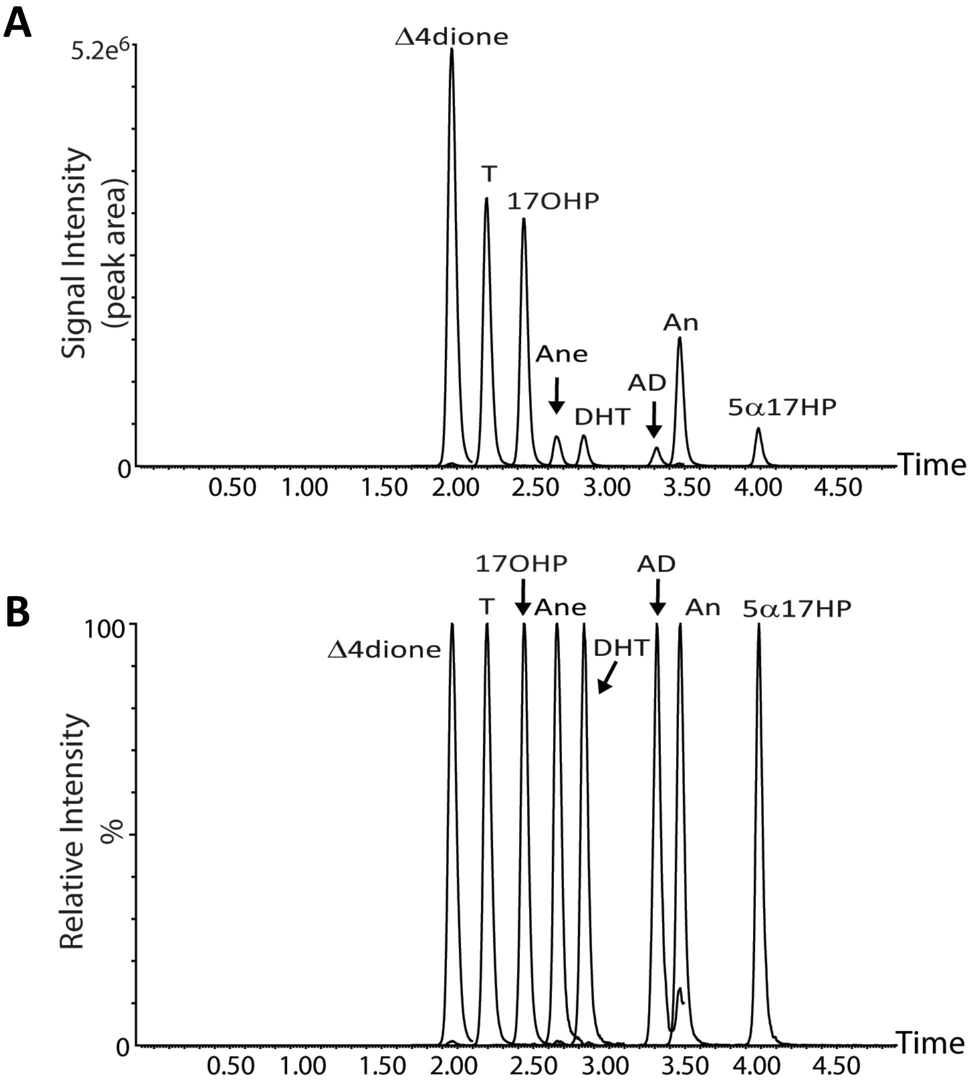


**Fig. S2. Representative liquid chromatography-tandem mass spectrometry (UHPLC-MS/MS) chromatograms**, produced on a Waters Xevo Mass Spectrometer in positive ionization mode, demonstrating optimized separation of mixture of authentic steroid standards. (**A**) Actual peak intensity (peak area), illustrating differences in ionization efficiency; (**B**) Relative peak intensity normalized to 100% for each standard.

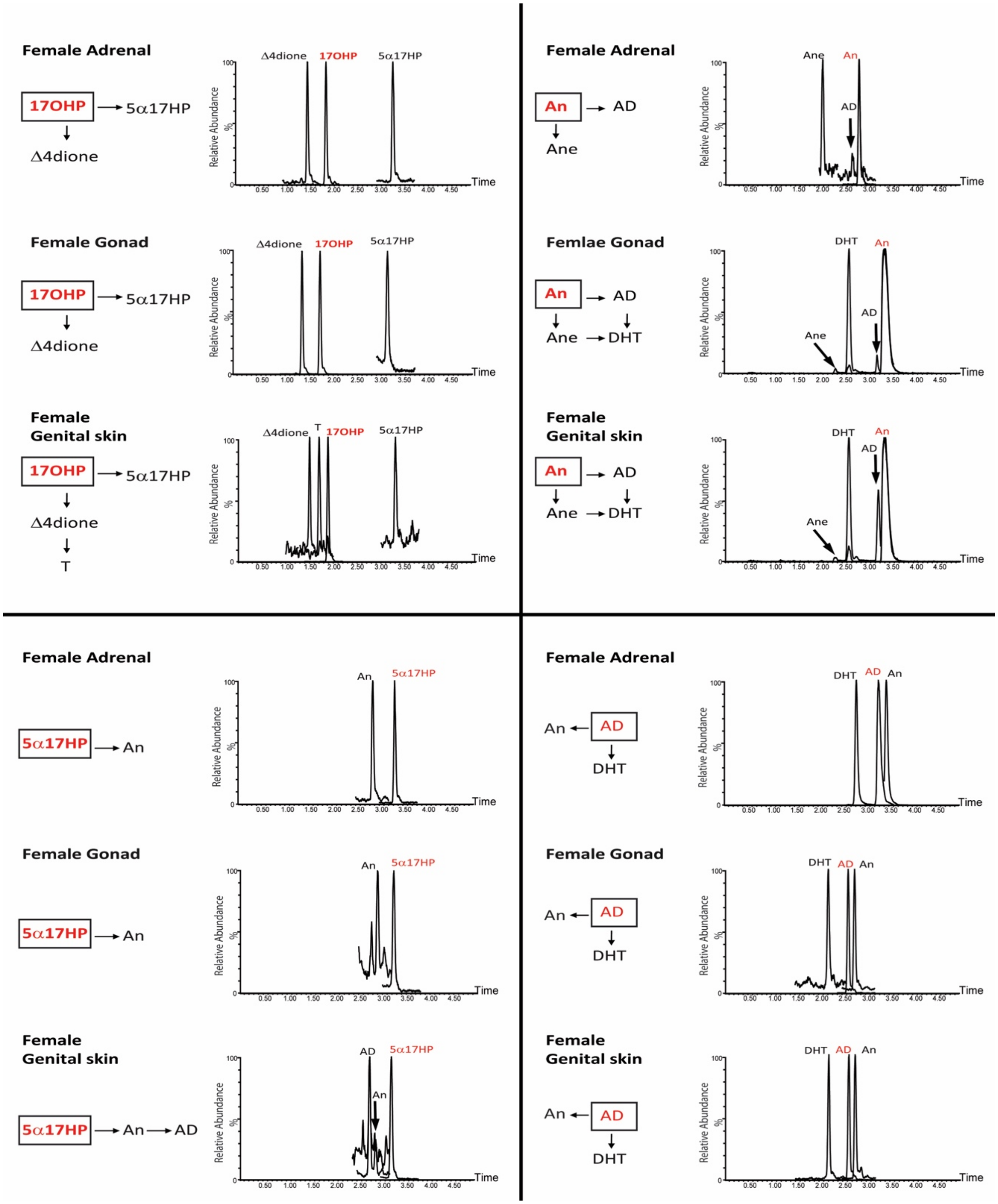


**Fig. S3. Steroid incubations of female fetal tissue explants**

The female adrenals, gonads and genital skin incubated with either (A) 17OHP, (B) 5α-17HP, (C) 5α-androsterone, or (D) 5α-androstanediol. Samples were analysed using an optimised LC-MS/MS method for qualitative identification of eight steroids; 17-hydroxyprogesterone (17OHP), 5α-17-hydroxypregnanolone (5α-17HP), androstenedione (Δ4dione), 5α-androsterone (An), 5α-androstanediol (AD), 5α-androstanedione (Ane), testosterone (T) and 5α-dihydrotestosterone (DHT). Substrates shown in red, products in black. Positive identification was based on matching retention time and two matching MRM transitions compared to the steroid standard. Representative chromatograms are shown for each incubation. Peaks represent the sum of the two mass transitions listed in **Table S1**.

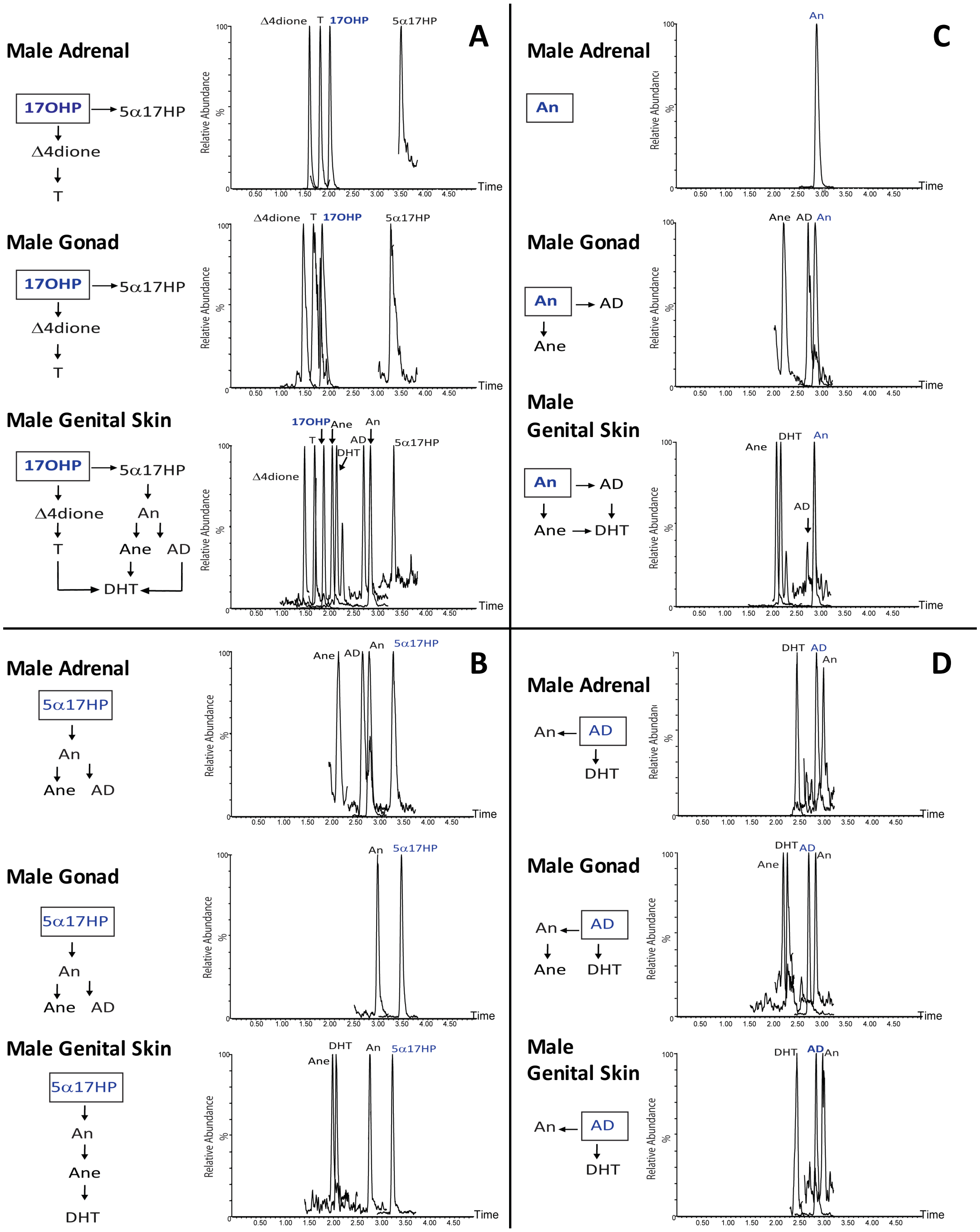


**Fig. S4. Steroid incubations of male fetal tissue explants**. The male adrenals, gonads and genital skin incubated with either (A) 17OHP, (B) 5α-17HP, (C) 5α-androsterone, or (D) 5α-androstanediol. Samples were analysed using an optimised LC-MS/MS method for qualitative identification of eight steroids; 17hydroxyprogesterone (17OHP), 5α-17-hydroxypregnanolone (5α-17HP), androstenedione (Δ4dione), 5α-androsterone (An), 5α-androstanediol (AD), 5α-androstanedione (Ane), testosterone (T) and 5α-dihydrotestosterone (DHT). Substrates shown in blue, products in black. Positive identification was based on matching retention time and two matching MRM transitions compared to the steroid standard. Representative chromatograms are shown for each incubation. Peaks represent the sum of the two mass transitions listed in **Table S1.**

**Table S1.** Retention times and mass transitions (quantifier and qualifier (italics) ions) of uPLC-MS/MS analysis of key steroids in the classic and alternative androgen biosynthesis pathways.

| **Steroid** | **Abbre-viation** | **Chemical name** | **Retention Time (min)** | **Mass Transitions (precursor>product)** |
|---|---|---|---|---|
| 17α-hydroxy-progesterone | 17OHP | 4-Pregnen-17α-ol-3,20-dione | 2.02 | 331 > 109<br>*331 > 97* |
| Androstenedione | $\Delta^4$dione | 4-Androstene-3,17-dione | 1.51 | 287 > 97<br>*287 > 109* |
| Testosterone | T | 4-Androsten-17β-ol-3-one | 1.71 | 289 > 97<br>*289 > 109* |
| 5α-dihydro-testosterone | DHT | 5α-Androstan-17β-ol-3-one | 2.41 | 291 > 255<br>*291 > 159* |
| 5α-17-hydroxy-pregnanolone | 5α-17HP | 5α-Pregnane-3α,17α-diol-20-one | 3.56 | 335 > 135<br>*335 > 299* |
| 5α-androsterone | An | 5α-Androstan-3α-ol-17-one | 3.04 | 291 > 273<br>*291 >255* |
| 5α-androstanediol | AD | 5α-Androstane-3α,17β-diol | 2.90 | 275 > 257<br>*275 > 147* |
| 5α-androstanedione | Ane | 5α-Androstane-3, 17-dione | 2.36 | 289 > 253<br>*289 > 213* |

**Table S2.** Metabolism of deuterated steroids by human fetal organ explant cultures (tissues collected from fetal material from the 6th to 10th week of gestation).

| Steroid | Abbre-viation | Retention Time (min) | Mass Transitions |
|---|---|---|---|
| **Precursor 17-hydroxyprogesterone-[2,2,4,6,6,21,21,21 $^2H_8$] (d8)** | | | |
| *Products Identified:* | | | |
| Androstenedione-d5 | $\Delta^4$dione-d5 | 1.51 | 292 > 100 |
| Testosterone-d5 | T-d5 | 1.71 | 294 > 100 |
| 5α-dihydrotestosterone-d5 | DHT-d5 | 2.41 | 296 > 278 |
| 5α-17-hydroxypregnanolone-d8 | 5α17HP-d8 | 3.56 | 343 > 140 |
| 5α-androsterone-d5 | An-d5 | 3.04 | 296 > 278 |
| **Precursor 5α-androsterone-[16,16 $^2H_5$ ] (d2)** | | | |
| *Products identified:* | | | |
| 5α-androstanediol-d2 | AD-d2 | 2.90 | 277 > 259 |
| 5α-androstanedione-d2 | Ane-d2 | 2.36 | 291 > 255 |
| 5α-dihydrotestosterone-d2 | DHT-d2 | 2.41 | 293 > 159 |
| **Precursor 5α-androstanediol-[16.16,17 $^2H_3$ ] (d3)** | | | |
| *Products identified:* | | | |
| 5α-androsterone–d2 | An-d2 | 3.04 | 293 > 275 |
| 5α-dihydrotestosterone-d3 | DHT-d3 | 2.41 | 294> 159 |
